# Influenza A virus H5N1 genotypes B3.13 and D1.1 show temperature-dependent restriction of replication in primary human respiratory epithelial cell cultures derived from the upper and lower respiratory tract

**DOI:** 10.64898/2026.08.27.747488

**Authors:** Anne P. Werner, Jaiprasath Sachithanandham, Elgin Akin, Sattya N. Talukdar, Matthew Pinsley, Andrew Pekosz

**Author notes:** Corresponding author: Andrew Pekosz.

## Abstract

H5N1 clade 2.3.4.4b avian influenza A viruses pose a significant threat to wild animal populations, domesticated animals, and potentially, the human population. For H5N1s to infect and transmit among mammalian species, mutations for improved utilization of mammalian receptors and enhanced replication at the lower temperatures of the upper respiratory tract need to be acquired. A human H1N1pdm09-like virus was compared to H5N1 genotypes B3.13 and D1.1 for replication at 33°C, 37°C, and 39°C – temperatures consistent with the upper and lower respiratory tract in humans, and dairy cow udder tissue. All H5N1 viruses had increased plaque sizes on MDCK cells at 37°C and 39°C compared to H1N1pdm09. In primary, differentiated human nasal and bronchial epithelial cultures, all H5N1 viruses show restricted infectious virus production compared to H1N1 at 33°C. While H5N1 D1.1 also showed restricted replication at 37°C and 39°C, the H5N1 B3.13 replicated to nearly equivalent titers as H1N1pdm09. All H5N1 viruses demonstrated similar cell tropism in cells from the upper and lower respiratory tract, infecting more ciliated than non-ciliated cells relative to H1N1pdm09. H1N1, H5N1 B3.13 D1.1 infection induced similar innate immune factors, with nasal epithelial cells producing higher levels compared to bronchial epithelial cells. These data suggest that genotype B3.13 and D1.1 H5N1 viruses show different temperature dependent replication patterns compared to H1N1pdm09.

**IMPORTANCE:** The ongoing H5N1 outbreaks have increased the number of mammalian infections with H5N1, thus increasing the chances of the virus adapting to mammals. Two H5N1 genotypes that have recently infected dairy cows, commercial poultry flocks, and a limited number of humans in the United States showed decreased replication at 33°C but not 37°C or 39°C compared to human H1N1 virus. H5N1 virus replication was detected at all temperatures, suggesting temperature and not host species may be the primary barrier preventing H5N1 infection in humans.

## INTRODUCTION

Highly pathogenic avian influenza (HPAI) viruses have been a major threat to the international poultry industry since its first detection in the 1980s ^1,2^. Primarily distinguished from their low-path (LPAI) counterparts by the presence of a multibasic cleavage site (MBS) in the hemagglutinin (HA) glycoprotein, HPAIs are characterized by a rapid decimation of domestic poultry ^3–5^. Substantiating additional concern surrounds their high morbidity and case fatality rates (CFRs) in humans during infrequent outbreaks and spillovers ^6–14^.

In 2024, H5N1 clade 2.3.4.4b moved into dairy cattle in the United States ^10,15–20^. H5N1 clade 2.3.4.4b genotype B3.13 viruses are responsible for a majority of cattle infections and for 41 of the 70 confirmed H5N1 infections in humans ^21^. A second H5N1 clade 2.3.4.4b genotype, D1.1, was the predominant genotype circulating among migratory birds, has been driving most commercial and backyard poultry outbreaks, has entered dairy cows on several separate occasions, and is the genotype responsible for two documented human fatalities in Louisiana and Mexico and a critical case in British Columbia, Canada ^11,22,23^. Genotypes D1.1 and B3.13 are both the progeny of distinct reassortment events between Eurasian lineage H5N1 viruses and LPAI viruses circulating in North America, likely contributing to their distinct circulation patterns and host ranges ^18,24^. The reassortments that yielded genotypes B3.13 and D1.1 further highlight the risk of co-infection in wild birds leading to the generation of genotypes that may have different replication, pathogenesis, and transmission profiles based on the specific gene constellation present ^25–31^.

Risk assessments of high consequence zoonotic influenza viruses rely on the integration of three main categories: public (i.e., human) health risks, animal health risks, and characterization of virus fitness in models of potential hosts ^32–34^. Previous work has suggested that ancestral H5N1 HPAI viruses demonstrate reduced growth in mammalian upper respiratory tract systems, largely contributable to poor binding of human-type sialic acid receptors and polymerase incompatibilities ^8,35–40^. While receptor binding profiles have been described for clade 2.3.4.4b H5N1s, there is limited data translating these findings into virus replication phenotypes in relevant, human respiratory cell culture systems and at physiologically relevant temperatures ^37,41–43^. This work investigates virus replication fitness, cell tropism, and induced innate immune responses in primary human respiratory epithelial cultures of two genotype B3.13 and one genotype D1.1 viruses, relative to a contemporary seasonal H1N1pdm09-like virus. Replication phenotypes and antiviral response states for the B3.13 and D1.1 viruses are largely influenced by temperature and cell type.

## MATERIALS & METHODS

### Viruses and virus stock preparation

H5N1 influenza virus isolates used in the study, A/Texas/37/2024 (GISAID Accession no. EPI_ISL_19027114), clade 2.3.4.4b, genotype B3.13; A/California/147/2024 (GISAID Accession no. EPI_ISL_19481947) clade 2.3.4.4b, genotype B3.13; and A/Washington/239/2024 (GISAID Accession no. EPI_ISL_19512045) clade 2.3.4.4b, genotype D1.1. All H5N1 isolates were kindly provided by the U.S. Centers for Disease Control (CDC). All H5N1 viruses were propagated on MDCKIs at 37°C and sequence-verified before use in assays as previously described ^44,45^. A/Baltimore/JH-640/2023 (H1N1pdm09-like) was isolated from the nasal swab of an infected patient presenting to Johns Hopkins Hospital (Baltimore, MD) during the 2023-2024 Northern Hemisphere Influenza season. Virus transport media containing the patient’s nasal swab was used to infect human nasal epithelial cell (hNEC) cultures, and the virus isolate was propagated on MDCK-SIAT1 cells as previously described ^45,46^.

### Biosafety Level Containment

All work with H5N1 viruses was performed at biosafety level 3 (BSL-3) using institution approved biosafety procedures. While seasonal H1N1 is a BSL-2 pathogen, all experiments were performed under BSL-3 conditions to maintain consistency with the H5N1 experiments.

### Immortalized cells

Madin-Darby Canine Kidney I (MDCKI) cells (Dr. Robert A. Lamb, Northwestern University) were used for generation of H5N1 stocks, plaque assays, and TCID_50_ assays. MDCKI cells stably over-expressing human sialyltransferase-1 (MDCK-SIAT1, Dr. Scott Hensley, University of Pennsylvania) were used to propagate seasonal H1N1pdm09-like stocks. Cells were maintained as previously described ^44–46^. Briefly, cells were incubated at 37°C/5% CO_2_ in complete medium (CM), comprised of Dulbecco’s Modified Eagle Medium (DMEM), supplemented with 10% fetal bovine serum (FBS, Gibco), 100 units/ml penicillin/streptomycin (Life Technologies), and 2 mM of GlutaMAX (Gibco).

### MDCKI Plaque assay

Plaque assays were performed on MDCKI cells as previously described ^47–49^. Briefly, MDCKI cells were seeded on 6-well tissue culture plates (Celltreat) two days prior to infection in CM. At the time of infection, media was removed and cells were washed twice with 1X PBS supplemented with Ca^2+^ and Mg^2+^ (PBS (+/+)), and 250 µL of virus inoculum was added to each well. Inoculum was incubated on cells for 1 hour, rocking, at 33°C, 37°C, or 39°C. Virus inoculum was then removed and replaced with phenol red-free MEM (Gibco) supplemented with 3% bovine serum albumin (Sigma-Aldrich), 100 units/ml penicillin/streptomycin (Life Technologies), 2 mM of GlutaMAX (Gibco), 5 µg/mL Trypsin acetylated from bovine pancreas (Sigma-Aldrich), 5 mM HEPES buffer (Gibco), and 1% cellulose was added. Cells were incubated at 33°C, 37°C, or 39°C for 60 hours. Cells were fixed with neutral buffered formalin (Leica) overnight and stained with naphthol blue-black (Sigma-Aldrich). Plaque area was quantified using ImageJ software as previously described ^47–49^.

### Primary cell cultures

Human nasal epithelial cells (PromoCell, Cat no. C-12620, Lot no. 482Z006.2) and human bronchial epithelial cells (Lonza; Cat no. CC-2540, Lot no. 0000613767) were grown on 6.5-mm transwells with 0.4-µm pore size (CellTreat). After commercial tubes were thawed, cells were expanded and transferred to collagen-coated transwells and maintained in PneumaCult™-ExPlus basal media (StemCell, Cat no. 05040). Transepithelial electrical resistance (TEER) was measured every other day until readings consistently exceeded 400 Ω•cm^2^, at which point apical media was removed and basal media was replaced with PneumaCult™-ALI basal medium (StemCell, Cat no. 05001). Cells were monitored daily to evaluate proper development of cilia and mucus production. All media was prepared per manufacturer’s instructions. All cells were grown at 37°C with 5% CO_2_ and 100% humidity prior to infection.

### Growth curves

All virus inoculum was prepared to an MOI of 0.01 by diluting virus in DMEM (Gibco). Wells were washed two times with 1X PBS+/+, and 100 µL of inoculum or DMEM alone was added to the apical side of transwells and incubated for 2 hours at 33°C/5% CO_2_, 37°C/5% CO_2_, or 39°C/5% CO_2_. After 2 hours, inoculum was removed and wells were washed two times with 1X PBS+/+. Plates were returned to incubators at one of three temperatures (33°C, 37°C, and 39°C) for a total duration of 120 hours. At each timepoint, 100 µL of DMEM (Gibco) was added to the apical side of each well, and plates were returned to their respective incubators for 10 minutes. The apical media was then removed and used to quantify infectious virus titer by TCID_50_. Basal media (PneumaCult™-ALI basal medium, StemCell) was replaced prior to infection and every 48 hours later.

### TCID_50_ assay

Tissue culture infectious dose 50% (TCID_50_) assays were used to quantify infectious virus in apical washes from hNEC and hBEC cultures as previously described ^45–48^. In brief, MDCKI cells were seeded in complete media on 96-well tissue culture plates (Celltreat) two days prior to infection. At the time of infection, wells were washed twice with PBS (+/+). Apical washes were serially diluted 10-fold, 8 times in infection media, which comprised of DMEM supplemented with 3% bovine serum albumin (Sigma-Aldrich), 100 units/ml penicillin/streptomycin (Life Technologies), 2 mM of GlutaMAX (Gibco), and 5 µg/mL Trypsin acetylated from bovine pancreas (Sigma-Aldrich). Diluted virus was added to cells in sextuplicate and incubated at 33°C/5% CO_2_ for 6 days. On the 6^th^ day post-infection, cells were fixed with neutral buffered formalin (Leica) for a minimum of 1 hour and subsequently stained with naphthol blue-black (Sigma-Aldrich) for further quantification.

### Confocal Microscopy

Differentiated hNECs and hBECs were infected at an MOI of 2 for two hours at 37°C. The inoculum was removed and cells were returned to the incubator for 24 hours. The transwells were washed with 1X PBS+/+, and both apical and basal sides of the transwells were exposed to 4% PFA in 1X PBS+/+ for 30 minutes at room temperature. Transwells were then washed an additional two times with 1X PBS+/+, followed by the addition of immunofluorescence (IF) buffer (130 mM NaCl_2_, 7 mM Na_2_HPO_4_, 3.5 mM NaH_2_PO_4_, 7.7 mM NaN_3_, 0.1% BSA, 0.2% Triton X-100, and 0.05% Tween 20), supplemented with 10% goat serum for 1 hour. Transwells were washed twice and then incubated with anti-influenza A virus nucleoprotein mouse monoclonal (HB65, 1:50 dilution) and Acetyl-α-tubulin rabbit monoclonal (1:500) (Cell Signaling Technologies, 5335), in IF washing buffer overnight at 4°C. Transwells were then washed and secondary antibodies, anti-rabbit Alexa Fluor 647 (1:200) (Thermo Fisher Scientific) and anti-mouse Alexa Fluor 488 (1:200) (Thermo Fisher Scientific), were diluted in IF buffer and added to transwells for 3 hours at 4°C. Transwells were then incubated with rhodamine phalloidin (PHDR1) (1:500) (Cytoskeleton, Inc.) for 30 min at 4°C, and nuclei were stained with NucBlue Fixed Cell Stain Ready probes (Thermo Fisher Scientific) for 30 min at 4°C. The transwell membrane was washed with IF buffer, and then cut and set on microscope slides (TechMed Services), and ProLong Gold antifade mounting medium (Life Technologies) was used for mounting coverslips. Images were captured using a confocal microscope (Leica Stellaris Falcon 8) enabled with a 63X objective. White light laser (WLL) and Power HyD detectors were used to find DAPI (49,6-diamidino-2-phenylindole) signal for nucleus detection, to activate Alexa Fluor 488 for Nucleoprotein detection, to activate rhodamine phalloidin for F-actin detection, and to activate Alexa Fluor 647 for Acetyl-a-tubulin detection. Imaris software version 10.1 (Oxford Instruments Group) was used for the conversion of z-stack images (.oir format) to.tiff format and other image post-processing.

### Image Analysis

The images were initially processed with Imaris software version 10.1 (Oxford Instruments Group) and used for the conversion of Z-stack images (.oir format) to.tiff format. Ciliated cells were detected by the presence of acetyl-α-tubulin (cyan) signal and non-ciliated cells were detected by the absence of acetyl-α-tubulin signal. Individual infected cells were confirmed by the presence of both F-actin (red) and nucleoprotein (NP) (green) signal (F-actin^+^/NP^+^). Individual infected cells with acetyl-α-tubulin signal (cyan) were considered infected ciliated cells, and individual infected cells without acetyl-α-tubulin signal were considered infected non-ciliated cells. At least 100 NP^+^ cells were selected randomly from both hNECs and hBECs for image analyses.

### Cytokine, chemokine, IFN-β and IFN-λ quantification

Basolateral media was inactivated by the addition of Tween-20 (Millipore Sigma) to a final concentration of 1% (v/v) and incubated for 10 minutes at ambient temperature before removal from BSL-3 containment. Secreted cytokines and chemokines were quantified using the 20-plex ProcartaPlex Human Inflammation Panel (ThermoFisher, EPX200-12185-901, lot# 462424-001) from basolateral media collected at 48 and 96 hours post-infection. The panel measured the following analytes: cytokines (GM-CSF, IFN-α, IFN-γ, IL-1α, IL-1β, IL-4, IL-6, IL-8, IL-10, IL-12p70, IL-13, IL-17A, TNF-α); chemokines (IP-10/CXCL10, MCP-1/CCL2, MIP-1α/CCL3, MIP-1β/CCL4); and adhesion molecules (sICAM-1, E-selectin/CD62E, P-selectin/CD62P). The assay was performed according to manufacturer’s instructions and acquired on the Luminex xMAP INTELLIFLEX System. Standard curves were generated and analyte concentrations (pg/mL) were calculated using the ProcartaPlex Analysis App (ThermoFisher). IFN-β was quantified by ELISA, per manufacturer’s instructions (PBL Assay Science, Cat no. 41435). IFN-λ was quantified by ELISA per manufacturer’s instructions (PBL Assay Science, Cat no. 61840). Data analysis and visualization were performed using custom R scripts with ggplot2, pheatmap, emmeans, and all base R packages.

## Data availability

All raw data is available at the Johns Hopkins Data Repository (DOI:XXXXXXXX. R scripts developed for Procartaplex analysis are available at https://github.com/[ANNIE TO FILL THIS IN!]

## RESULTS

### Genetic characterization of H5N1 clade 2.3.4.4b isolates

Three clade 2.3.4.4b isolates from confirmed human infections were selected for investigation: A/Texas/37/2024 and A/California/147/2024 (genotype B3.13) and A/Washington/239/2024 (genotype D1.1). The two B3.13 viruses were obtained from dairy farmworkers that both had worked with H5N1-infected cows ^12,50^, and the D1.1 virus was obtained from a poultry farmworker who had known exposures to infected birds ^51^. A H1N1 pandemic 2009 (H1N1pdm09)-like isolate—A/Baltimore/JH-640/2023—was obtained from Johns Hopkins Hospital (JHH) seasonal surveillance during the 2023-2024 Northern Hemisphere (NH) season.

Multiple sequence alignments (MSA) of the polymerase basic 2 (PB2) gene, the hemagglutinin (HA) gene, and the neuraminidase (NA) gene (**Table 1**), as well as the remaining 5 segments (**Table S1**) were performed. The alignment revealed some heterogeneity within the B3.13 genotype (**Table 1**)., There are a considerable number of amino acid differences in all 8 segments between B3.13 genotype isolates and the D1.1 isolate (**Table 1 & S1**), which was expected as the two genotypes are derived from distinct 4:4 reassortment events ^18^. Although only A/Texas/37/2024 contained the E627K mutation for mammalian adaptation in PB2, A/California/147/2024 contained M631L, another mutation known to increase avian virus replication in mammalian tissues ^39,52–54^. Other known PB2 mutations associated with enhanced replication in human respiratory epithelium, i.e., E249G, T339M, D701N, were not present in any of the three H5N1 isolates (**Table 1**) ^55^.

**Table 1.** Amino acid substitutions in A/H5N1 2.3.4.4b isolates (relative to A/Texas/37/2024). All HA sequences shared the following residues at positions known to influence transmission and/or virulence: 110H, 119E, 152V, 158N, 160I, 224N, 226Q, 228G, 318T

| Isolate name | Genotype | Segment name |  |  |
| --- | --- | --- | --- | --- |
|  |  | PB2 | HA (H3 numbering) | NA |
| A/Texas/37/2024 | B3.13 | -- | -- | -- |
| A/California/147/2024 | B3.13 | A58T, E362G, K627E, M631L, K670R | D95S, **V135M**, S323N | N71S |
| A/Washington/239/2024 | D1.1 | A58T, I109V, I139V, N441D, I495V, K627E | T46A, **M111L**, **Q122L**, **I199T**, **A214V**, 326R*, N475D, V510I | T8I, V20I, M23V, Y44N, Q45H, P48T, I53V, F74L, L75I, T81D, S82P, T84A, N220S, V234I, V241I, K257R, M269L, G286S, D287E, I288V, I321V, N329S, S336G, M338V, P339S, E395A, K432R |

\*furin cleavage site absent in H3, no numbering available

The HA was largely conserved across genotypes with the exception of several amino acid substitutions within defined antigenic regions and the receptor binding domain (RBD) (**Table 1**, shown in bolded and red bolded font, respectively). Importantly, there were no detected amino acid mutations that are known to increase virulence in mammals nor were there mutations that conferred ɑ2,6-linked SIA-binding ^35,36,41,56,57^. Only A/Washington/239/2024 contained a mutation in the RBD ^18,58^.

The NA gene of A/Washington/239/2024 was the most divergent segment between the three H5N1 viruses, due to the fact it was derived from a distinct LPAI lineage than that in genotype B3.13 viruses ^18^. Twenty seven unique NA amino acid mutations were detected between A/Washington/239/2024 and A/Texas/37/2024 as well as A/California/147/2024 (**Table 1**), with many near the catalytic site (N220S), secondary sialic acid binding sites (E395A and K432R) and antigenic sites I, II, IV, and V (**Table 1**) ^59,60^. The mutations in the NA segment were distinct in A/Washington/239/2024 but conserved in both B3.13 viruses (**Table 1**).

The remaining polymerase segments, PB1 and PA, of A/Washington/239/2024, in addition to the NA segment, showed a number of differences (**Table S1**). Both A/Washington/239/2024 and A/California/147/2024 shared PB1 V392I and PA E142K distinct from A/Texas/37/2024. Recent studies investigating pathogenesis and virulence of a reverse genetics virus containing the PA E142K mutation suggests that this mutation may interact with pathways involved in CCL2-mediated signaling and mitigate severe disease outcomes in mouse models ^61^.

NS1 proteins from A/Texas/37/2024, A/California/147/2024, and A/Washington/239/2024 showed limited amino acid variation across the RNA-binding, linker, and effector domains ^62^ (**Table S1**). Compared with A/Texas/37/2024, A/California/147/2024 differed at only two RNA-binding domain residues, R40Q and R67G. R40Q has been observed in bovine H5N1 isolates but remains functionally uncharacterized, whereas residue 67 has been implicated in interactions with host translation factors and NS1 thermostability ^63^. A/Washington/239/2024 was more divergent, encoding substitutions in the RNA-binding domain (L7S, R40Q), linker region (E75G, S83P, P87S), and effector domain (S116C, N139D, L147I) ^64^. L7S has been reported during avian-to-mammalian spillover events, although available data indicate minimal effect on IFN-β promoter activity *in vitro* ^64^. S83P, proximal to the linker– effector domain interface, may potentially alter local flexibility or phosphorylation-dependent effects ^65,66^.

### H5N1 plaque morphology suggests a distinct temperature-sensitive phenotype that opposes seasonal H1N1 viruses

Plaque assays were performed in MDCKI cells at three distinct temperatures; 33°C to represent the temperature of the human upper respiratory tract, 37°C to represent human core body temperature, and 39°C to emulate the temperature of dairy cow udder tissue ^67^. The H1N1pdm09-like virus showed the largest plaque sizes at 37°C, which were only marginally larger than those at 33°C (**Fig. 1a**). In accordance with previously published work ^68,69^, the H1N1pdm09-like virus showed a marked reduction in plaque size at 39°C relative to 37°C and to 33°C (p < 0.0001 and p = 0.0013) (**Fig. 1a**). The largest plaque sizes observed among H1N1-infected cells were still smaller than plaques grown at 33°C for both genotype B3.13 H5N1 viruses (**Fig. 1b-c**).

**Figure 1.**
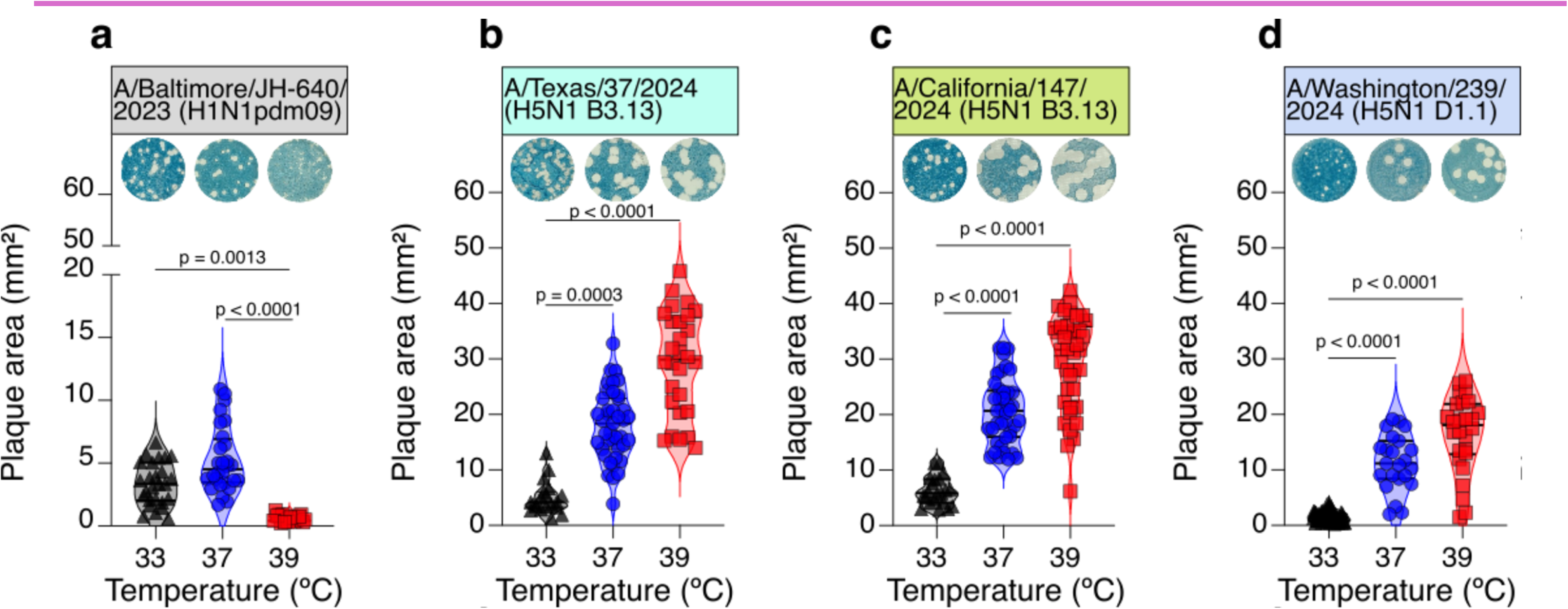
A/H5N1 viruses produce smaller plaques as temperature decreases. (**a-d**) MDCKI cells were infected with (**a**) a contemporary human seasonal A/H1N1 virus, (**b-c**) A/H5N1 genotype B3.13 viruses, or (**d**) an A/H5N1 genotype D1.1 virus, and incubated at 33°C, 37°C, or 39°C for 60 hours to evaluate differences in plaque morphology. (**a-d**) Representative images of each virus and temperature combination are shown. Each symbol represents the quantification of an individual plaque. Plaques were quantified in ImageJ. Data is combined from two or three independent experiments. P-values less than 0.05 are shown, as calculated via one-way ANOVA with Kruskal-Wallis nonparametric post-test for multiple comparisons between groups.

All three H5N1 isolates showed greater plaque area at higher temperatures (**Fig. 1b-d**) 33°C (**Fig. 1b-d**). Plaques formed by the genotype D1.1 virus, A/Washington/239/2024, were smaller than those formed by the two genotype B3.13 viruses at each temperature. Additionally, the average plaque size for A/Washington/239/2024 at 33°C was smaller than the average plaque size formed by A/Baltimore/JH-640/2023 at 33°C (1.62 mm^2^ and 3.35 mm^2^, respectively), suggesting that genotype D1.1 viruses have an attenuated phenotype at 33°C (**Fig. 1a & 1d**) when compared to B3.13 and H1N1pdm09-like viruses.

### H5N1 virus replication kinetics in the upper and lower respiratory tract epithelial cell cultures are enhanced by higher temperatures

Infectious virus production was quantified in both normal human nasal epithelial cell (hNECs) cultures and normal human bronchial epithelial cell (hBECs) cultures, which have been reported to have greater expression of ɑ2,6-linked SIAs and ɑ2,3-linked SIAs, respectively ^70–73^, after infection at an MOI of 0.01 (**Fig. 2 & 3**).

**Figure 2.**
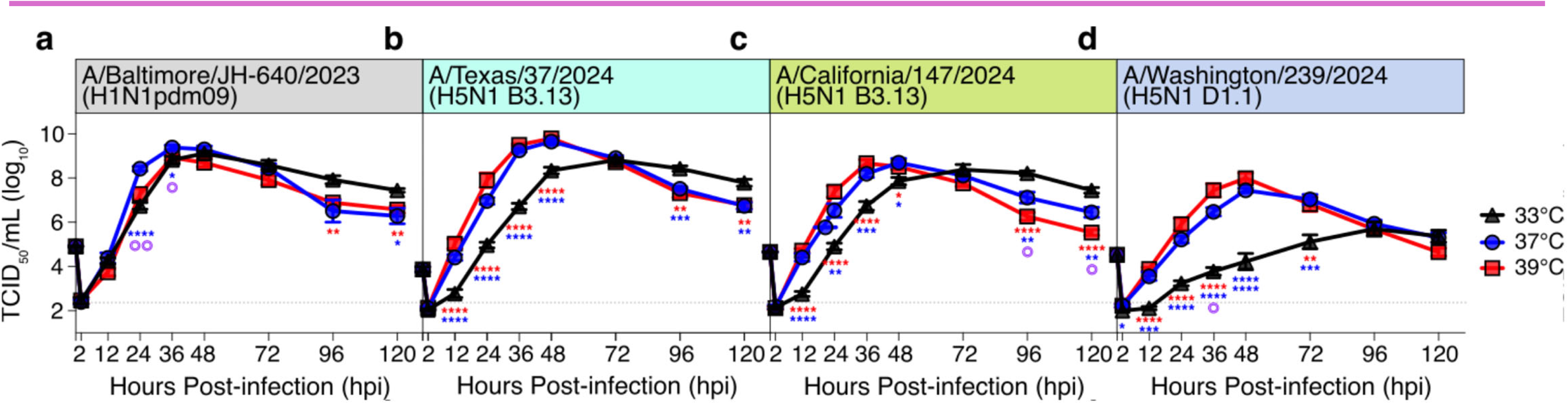
A/H5N1viruses show decreased infectious virus production at lower temperatures in human nasal epithelial cell cultures. (**a-d**) differentiated primary human nasal epithelial cells were grown in Transwells at an air-liquid interface (ALI), then infected at an MOI of 0.01 at 33°C (black triangles), 37°C (blue circles), or 39°C (red squares) with (**a**) a contemporary human seasonal H1N1 isolate, (**b-c**) A/H5N1 genotype B3.13 viruses, or (**d**) an A/H5N1 genotype D1.1 virus. (**a-d**). Dotted line represents the lower limit of detection of the TCID_50_ assay, and symbols and error bars represents the average and SEM of 8 individual replicates for each virus performed across two independent experiments. Asterisks (*) indicate significant differences between either 37°C and 33°C (indicated by blue asterisks), or 39°C and 33°C (indicated by red asterisks). (**a-d**) Purple bullseyes (◎) indicate significant differences between 37°C and 39°C. Significance was determined via two-way ANOVA with multiple comparisons between each of the three tested temperatures per timepoint.

In hNEC cultures, the H1N1pdm09-like virus replicated equally well across the three temperatures (**Fig. 2a**), whereas all H5N1 viruses showed lower peak infectious virus titers and slower kinetics of infectious virus production at 33°C than at 37°C or 39°C, indicating attenuation at lower temperatures (**Fig. 2b-d**). At 33°C, neither A/Texas/37/2024 B3.13 (**Fig. 2b**) nor A/Washington/239/2024 D1.1 (**Fig. 2d**) reached the same peak titers achieved at 37°C nor at 39°C (**Fig. 2b & 2d**). Only A/California/147/2024 B3.13 showed comparable peak virus titers during the course of infection in hNECs at 33°C relative to 37°C and 39°C (**Fig. 2c**) but this was reached 24-48 hours later. These temperature-dependent differences can be seen clearly when the data is grouped by the temperature of the experiment (**Fig. S1**). Across all temperatures, the D1.1 genotype virus showed reduced infectious virus production compared to the two B3.13 viruses and H1N1pdm09-like. At 33°C, the H1N1pdm09-like virus replicated to higher infectious virus titers and with faster kinetics than any of the H5N1 viruses (**Fig. S1a**) but at 37°C or 39°C, infectious virus production by B3.13 was similar to or even exceeded that of H1N1pdm09-like virus (**Fig. S1b-c**).This data demonstrates that B3.13 H5N1 viruses show slower kinetics of virus production at 33°C and produce comparable amounts of infectious virus as H1N1pdm09-like viruses over the entire course of the infection at all temperatures. In contrast, the H5N1 D1.1 genotype virus shows both slower kinetics and reduced total amounts of infectious virus compared to H5N1 B3.13 and H1N1pdm09-like viruses across all temperatures.

In hBEC cultures, the H1N1pdm09-like virus demonstrated similar kinetics and similar peak infectious virus titers across all three temperatures (**Fig. 3a**) while all three H5N1 viruses showed lower infectious virus titers at 33°C than at 37°C or 39°C, particularly at early timepoints (**Fig. 3b-d**). The genotype D1.1 virus replicates less effectively compared to both the H1N1pdm09-like and the H5N1 B3.13 viruses, regardless of temperature (**Fig. S2a-c**). Both genotype B3.13 viruses showed peak infectious virus titers at 36 HPI when incubated at 37°C or 39°C, which is similar to H1N1pdm09-like viruses, but incubation at 33°C delayed peak virus titers by 36 hours (**Fig. 3b-c**, **Fig. S2a**). The genotype D1.1 virus had peak titers that were 10^3^ and 10^4^-fold lower than peak titers at 37°C and 39°C, respectively (**Fig. 3c**, **Fig. S2a-c**). Despite previous literature implicating the higher relative expression of ɑ2,3-linked SIAs in the lower respiratory tract in enhanced H5N1 replication in hBECs ^42,73–75^, comparable replicative capacities were detected between the two cell culture systems with temperature rather than respiratory epithelial cell source being the variable most affecting infectious virus production.

**Figure 3.**
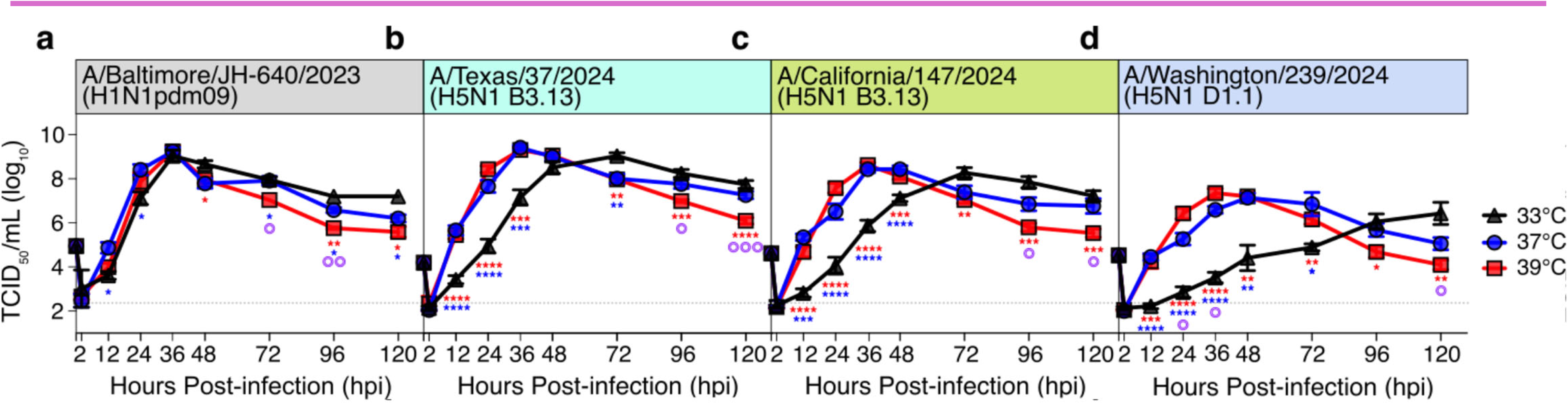
A/H5N1viruses show decreased infectious virus production at lower temperatures in human bronchial epithelial cell cultures. (**a-d**) differentiated primary human bronchial epithelial cells were grown in Transwells at an air-liquid interface (ALI), then infected at an MOI of 0.01 at 33°C (black triangles), 37°C (blue circles), or 39°C (red squares) with (**a**) a contemporary human seasonal H1N1 isolate, (**b-c**) A/H5N1 genotype B3.13 viruses, or (**d**) an A/H5N1 genotype D1.1 virus. (**a-d**). Dotted line represents the lower limit of detection of the TCID_50_ assay, and symbols and error bars represents the average and SEM of 8 individual replicates for each virus performed across two independent experiments. Asterisks (*) indicate significant differences between either 37°C and 33°C (indicated by blue asterisks), or 39°C and 33°C (indicated by red asterisks). (**a-d**) Purple bullseyes (◎) indicate significant differences between 37°C and 39°C. Significance was determined via two-way ANOVA with multiple comparisons between each of the three tested temperatures per timepoint.

### Cellular tropism of H1N1 and H5N1 viruses in upper and lower airway cultures

Differentiated hNECs and hBECs were infected at an MOI = 2 for 24 hours at 37°C by either H5N1 or H1N1pdm09-like viruses and acetyl-α-tubulin, a prominent marker of cilia, was used to distinguish ciliated from non-ciliated epithelial cells (**Fig. 4a & 5a**) ^76^. F-actin and NP were used as markers of cellular boundaries and viral antigen, respectively. In hNEC cultures, both H5N1 and H1N1pdm09 virus antigen was detected in ciliated and non-ciliated cells, with a preference for infection of ciliated cells across all viruses (**Fig. 4a-b**). Similar preferences for ciliated cells over non-ciliated cells were detected in hBEC cultures for all three H5N1 isolates, however, the H1N1pdm09-like virus infected significantly fewer ciliated cells than any of the H5N1 viruses (**Fig. 5a-b**). Taken together, the data indicate that H5N1 viruses prefer ciliated cells in both hNEC and hBEC cultures, while H1N1pdm09-like seasonal viruses have a reduced tropism for ciliated cells in hBEC cultures.

**Figure 4.**
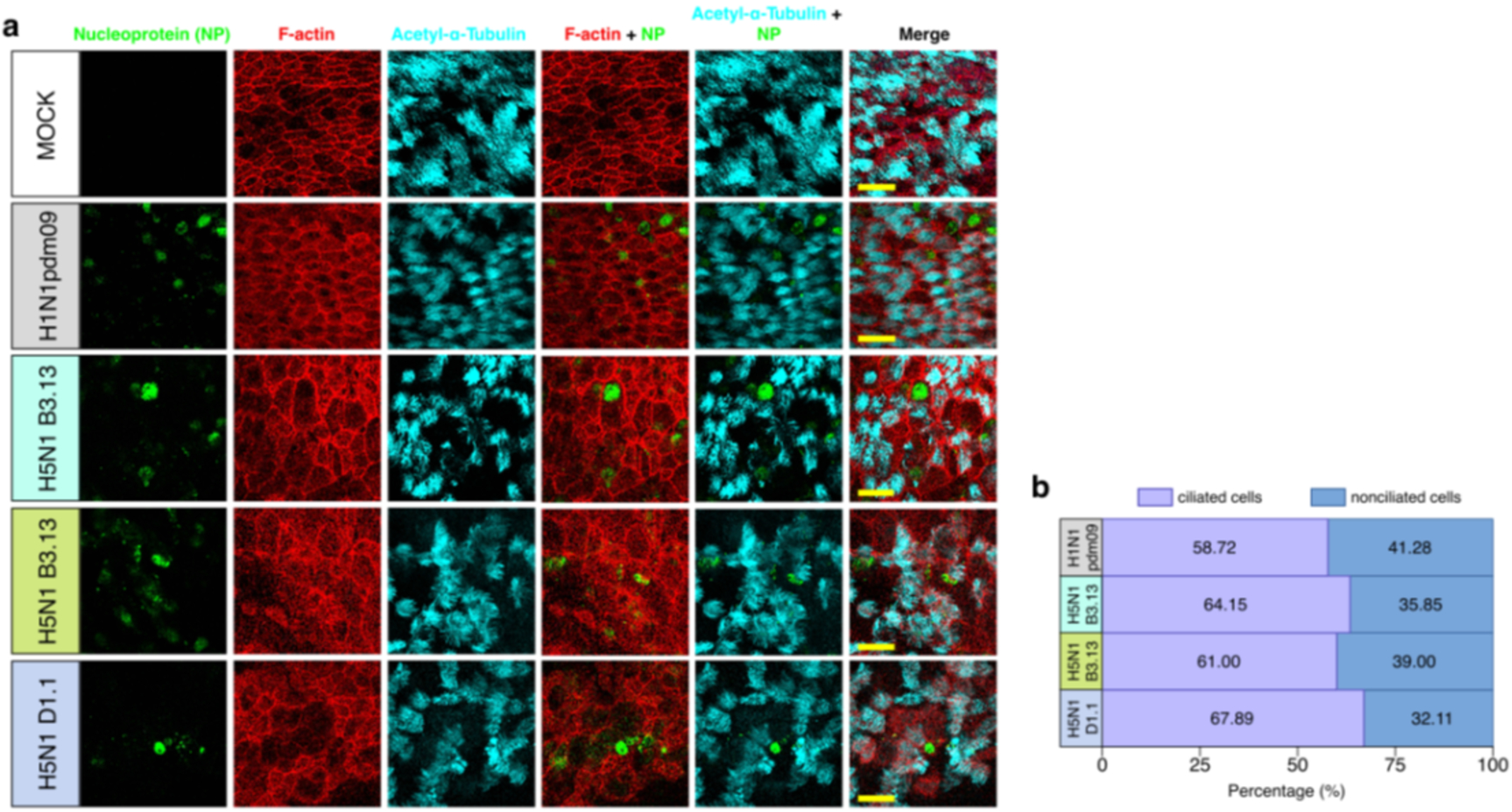
Infection of A/H5N1 viruses in differentiated human nasal epithelial cells, **(a)** Differentiated primary human nasal epithelial cells were grown in Transwells at air-liquid interface (ALI) culture, then infected at an MOI of 2 at 37°C for 24 hours. Cells were infected with a human seasonal H1N1pdmO9 isolate (A/Baltimore/JH-640/2023), two H5N1 genotype B3.13 viruses (A/Texas/37/2024 and A/California/147/2024) or one H5N1 genotype D1.1 virus (A/Washington/239/2024). Twenty-four hours post-infection, cells were fixed, permeabilized, and immuno-stained for nucleoprotein (green) and Acetyl-a-tubulin (cyan). F-actin was visualized by rhodamine phalloidin (red) staining. All images were captured with a 63X objective. Scale bar (Yellow) = 20 pm. **(b)** Ciliated and non-ciliated cell tropism during infection was quantified by I MARIS 10.1 software. At least 100 virus infected cells were randomly selected, then assessed for cell type infected.

**Figure 5.**
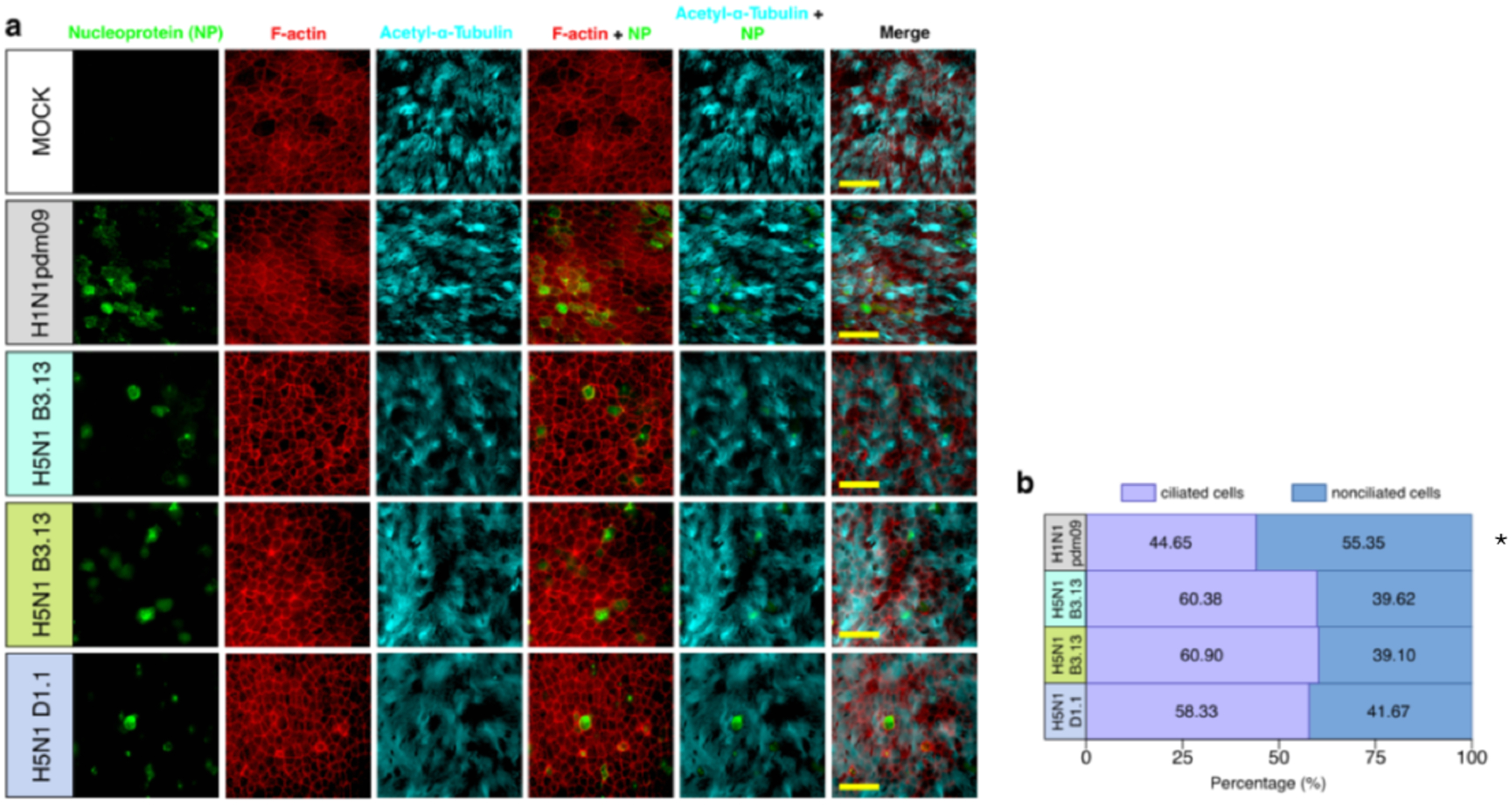
Infection of A/H5N1 viruses in differentiated human bronchial epithelial cells, **(a)** Differentiated primary human bronchial epithelial cells were grown in Transwells at air-liquid interface (ALI) culture, then infected at an MOI of 2 at 37°C for 24 hours. Cells were infected with a human seasonal H1N1pdmO9 isolate (A/Baltimore/JH-640/2023), two H5N1 genotype B3.13 viruses (A/Texas/37/2024 and A/California/147/2024) or one H5N1 genotype D1.1 virus (A/Washington/239/2024). Twenty-four hours post-infection, cells were fixed, permeabilized, and immuno-stained for nucleoprotein (green) and Acetyl-a-tubulin (cyan). F-actin was visualized by rhodamine phalloidin (red) staining. All images were captured with a 63X objective. Scale bar (Yellow) = 20 pm**. (b)** Ciliated and non-ciliated cell tropism during infection was quantified by I MARIS 10.1 software. At least 100 virus infected cells were randomly selected, then assessed for cell type infected. Statistical differences were determined by Chi-squared test (asterisk).

### The production of innate immune factors is influenced by respiratory epithelial cell type and temperature more than by virus type

Basolateral media was collected during H5N1 and H1N1pdm09-like virus infection and type I and III interferons (IFNs), and an additional 20 targets that are part of the canonical pro-inflammatory response were quantified at 48 (**Fig. 6**) and 96 HPI (**Fig. 7**). Hierarchical clustering revealed that relative quantities of interferons and pro-inflammatory signaling molecules clustered primarily by temperature and cell type, rather than by virus subtype or genotype (**Fig. 6-7**). At both timepoints, hBEC cultures clustered together, with most factors being downregulated or unchanged in production while hNEC cultures clustered together with somewhat higher expression levels (**Fig. 6-7**). At 48 HPI, there was also a clear effect of temperature on the production of factors such as CXCL10 (IP-10) and IL-8 in both culture types (**Fig. 6**), with both factors clustering together as downregulated at 33°C but having increased expression at higher temperatures. These temperature effects were not as obvious at 96 HPI (**Fig. 7**). A/Texas/37/2024 tended to cluster with seasonal H1N1pdm09-like virus instead of with the other two H5N1 isolates at 96 HPI (**Fig. 7**), rather than the general clustering of all H5N1 viruses at 48 HPI (**Fig. 6**).

**Figure 6.**
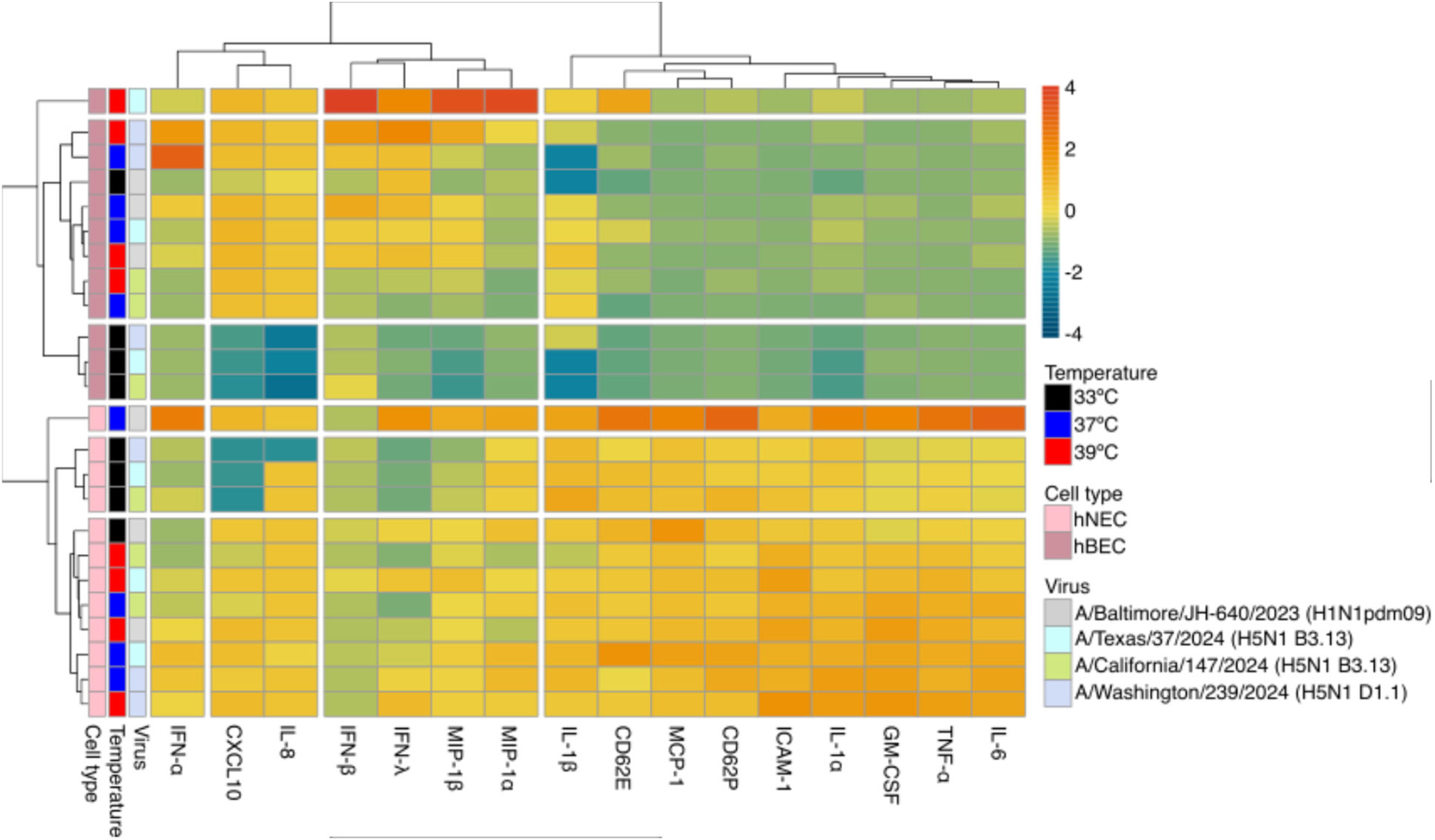
Chemokine and cytokine profiles in hNEC and hBEC cultures at 48 hours post-infection. Heat map showing relative quantity of the indicated analyte (IL-8, CXCL10, MIP-1ɑ, MIP-1β, IFN-β, IFN-λ, IL-1β, ICAM-1, IL-1ɑ, GM-CSF, IL-6, TNF-ɑ, CD62E, CD62P, MCP-1, IFN-ɑ, IL-10) as indicated by the scale on the right. Each tile represents the z-score (i.e., number of standard deviations away from the arithmetic mean) of the average concentration across 8 biological replicates (n = 4 per experiment) for the indicated analyte on the x-axis. Heatmap generated using the pheatmap package in RStudio.

**Figure 7.**
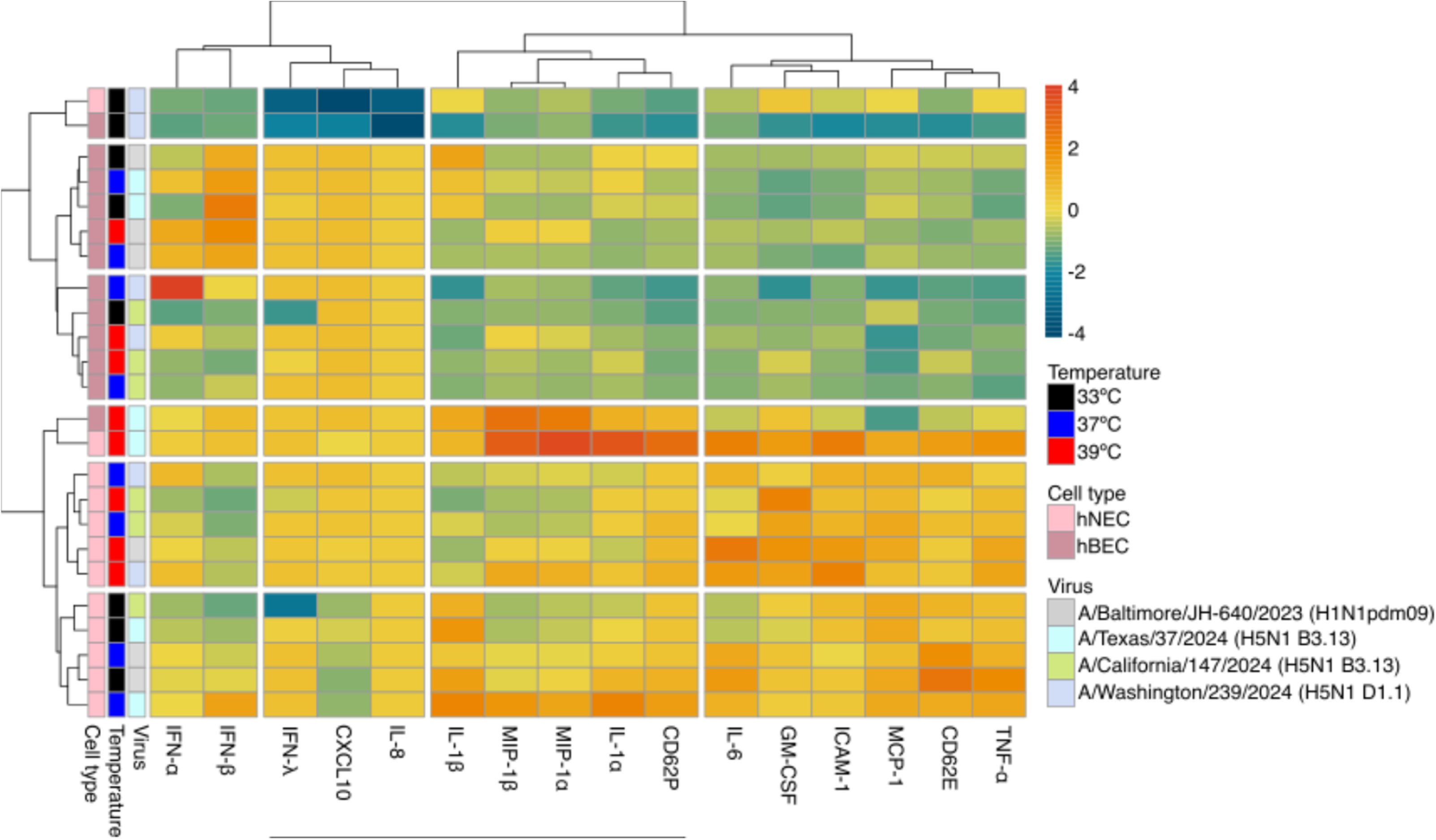
Chemokine and cytokine profiles in hNEC and hBEC cultures at 96 hours post-infection. Heat map showing relative quantity of the indicated analyte (IL-8, CXCL10, MIP-1ɑ, MIP-1β, IFN-β, IFN-λ, IL-1β, ICAM-1, IL-1ɑ, GM-CSF, IL-6, TNF-ɑ, CD62E, CD62P, MCP-1, IFN-ɑ, IL-10) as indicated by the scale on the right. Each tile represents the z-score (i.e., number of standard deviations away from the arithmetic mean) of the average concentration across 6-8 biological replicates (n = 2-4 per experiment) for the indicated analyte on the x-axis. Heatmap generated using the pheatmap package in RStudio.

At 48 HPI (**Fig. 6**) there was a marked reduction in quantity of signaling molecules IL-8 ^77–80^ and CXCL10 (IP-10) ^81–84^– which are critical for the recruitment of immune effector cells to the site of infection– for all three H5N1 viruses compared to the seasonal H1N1pdm09-like virus in both hNECs and hBECs (**Fig. 6**), but this difference was most pronounced in cells incubated at 33°C (**Fig. 6**). Levels of secreted CXCL10 in both cell types were significantly lower in cultures infected with any of the H5N1 isolates compared to cultures infected with the H1N1pdm09-like virus at 33°C (p < 0.001, **2, Fig. 6**). Additionally, A/California/147/2024-infected hBECs at 33°C had significantly lower quantities of CXCL10 than those infected with A/Texas/37/2024 and A/Washington/239/2024 (p < 0.001, **Table S2, Fig. 6**). H5N1-infected hNECs and hBECs showed significantly lower quantities of secreted IFN-λ at 33°C (p < 0.01 for A/Texas/37/2024, p < 0.001 for A/California/147/2024 and A/Washington/239/2024, **Table S2, Fig. 6**), and there was an observed reduction in type I interferons IFN-ɑ and IFN-β (**Fig. 6**). MCP-1 was significantly reduced in hNECs infected with any of the H5N1 isolates relative to the seasonal H1N1pdm09 isolate (p < 0.001, **Table S2, Fig. 6**). Regardless of infecting virus or temperature, hBECs typically secreted lower quantities of ICAM-1, IL-1ɑ, GM-CSF, IL-6, E-and P-selectins, and MCP-1 than hNECs, suggesting that the upper and lower respiratory tracts are poised differently to respond to H5N1 or H1N1 infection (**Fig. 6**).

Similar clustering of cytokines was observed at both timepoints, with CXCL10, IL-8, and type I IFNs clustering closely at both 48 HPI (**Fig. 6**) and 96 HPI (**Fig. 7**). Only A/Washington/239/2024-infected hNECs and hBECs at 33°C maintained low quantities of CXCL10 and IL-8 at 96 HPI, whereas comparable quantities of CXCL10 and IL-8 were observed between all other H5N1-and H1N1pdm09-infected cultures by 96 HPI (**Fig. 7**). Quantities of IFN-β remained low only for hNECs and hBECs infected with either the genotype D1.1 H5N1 virus, A/Washington/239/2024, or A/California/147/2024, relative to cultures infected with A/Texas/37/2024 or A/Baltimore/JH-640/2023, regardless of temperature (**Fig. 7**). Despite both being genotype B3.13 viruses, A/Texas/37/2024 induced significantly greater production of MIP-1β and MIP-1ɑ, in addition to IFN-β in hNECs and hBECs at 37°C and 39°C, but not at 33°C (**Table S3**, **Fig. 7**). The same trend was detected in A/Washington/239/2024-infected cultures relative to A/Texas/37/2024, such that significantly lower quantities of MIP-1β and MIP-1ɑ were detected but only at the two higher temperatures (**Table S3**, **Fig. 7**). In hBECs, both A/California/147/2024 and A/Washington/239/2024 induced significantly lower secretion of IL-1β at all three temperatures relative to A/Texas/37/2024. Altogether, these data show distinct cytokine and chemokine profiles influenced primarily by cell type and temperature but with some virus specific variation both across and within subtypes.

## DISUSSION

Evaluations of virus fitness and replication capacity in human primary cell cultures remains an essential component for risk assessment tools, like the World Health Organization’s Tool for Influenza Pandemic Risk Assessment (TIPRA) and the Centers for Disease Control and Prevention’s Influenza Risk Assessment Tool (IRAT) ^32–34^. Our data indicates that all three H5N1 isolates show reduced plaque size at 33°C– a temperature consistent with that of the human upper respiratory tract– compared to plaque sizes at 37°C, representing core human body temperature, or 39°C which approximates not only the internal temperature of udder tissue in dairy cattle ^67^ but also pyrexia in humans ^85^ (**Fig. 1a-d**). Furthermore, 39°C is closer to the internal or core temperatures of many avian species ^86^. The similar temperature restrictions on virus replication in air-liquid interface (ALI) hNEC and hBEC cultures (**Fig. 2-3, S1-3**) and plaque size phenotypes suggests that plaque size roughly correlates with overall infectious virus production in primary human respiratory epithelial cell cultures. The significant amounts of infectious virus produced from hNEC and hBEC cultures by H5N1 viruses indicates that the adaptions needed for virus replication in human respiratory cells are already present. However, in both hNEC and hBEC cultures, infectious virus production was more obviously reduced at early timepoints, suggesting that the efficiency of infection or the kinetics of infectious virus production were reduced for all three H5N1s at 33°C. Only the D1.1 H5N1 virus, A/Washington/239/2024, showed significant attenuation in total virus replication capacity in both hNEC and hBEC cultures relative to both B3.13 H5N1 viruses and the H1N1pdm09-like virus, and had consistently smaller plaque sizes than both B3.13 H5N1 viruses at all three temperatures.

Despite the clearly attenuated phenotype of D1.1 H5N1 replication in primary epithelial cultures, the D1.1 genotype isolate was able to produce plaques in MDCKIs that were larger than plaques formed by the H1N1pdm09-like virus at both 37°C and 39°C (p = 0.0003 & p < 0.0001, respectively), suggesting that adaptations needed for efficient replication in mammalian cells do not perfectly translate between immortalized and primary respiratory epithelial cell cultures. The observed reduction in replication fitness in the D1.1 genotype virus could be attributed to several things, including the availability of ɑ2,3-linked SIAs, and the susceptibility to innate antiviral responses mounted in the primary culture models. Another factor that could account for the differences observed in the plaque assay readouts and the hNEC and hBEC infectious virus production assays are differences in genome segments that are important for infectivity or in antagonizing certain innate signaling molecules. The major differences detected in the NA segment, which is critical for penetrating through mucus in the airway epithelium and effective cleavage of decoy receptors from HA ^87–91^, may partially explain the attenuation in epithelial cultures that is less pronounced in plaque assays. Other genetic differences observed in NP between the B3.13 and D1.1 genotype H5N1 may also contribute to enhanced susceptibility to Mx proteins, specifically H52Y ^39,92–95^. NP-52Y has also been reported to be critical in the interaction between host BTN3A3 and NP, such that 52Y confers sensitivity to BTN3A3-mediated antiviral signaling, resulting in markedly reduced virus replication *in vitro* ^95^. Expression of BTN3A3 throughout upper and lower respiratory epithelium ^96^, as well as constitutive expression in immortalized hBECs ^95^ indicate that these NP-host interactions may partially be responsible for the marked attenuation in D1.1 replication in hNECs and hBECs. This highlights a path of further study for characterizing primary hNEC and hBEC ALI culture systems and the expression of putative interacting host factors with avian and seasonal influenza viruses.

All three H5N1 isolates showed greater infection of ciliated cells versus non-ciliated cells in hNECs and hBECs. In contrast, seasonal H1N1pdm09-like virus infected fewer ciliated cells in hBEC cultures compared to the H5N1 viruses. Previous work has shown a bias toward ciliated cells in bronchial and nasal epithelial cultures for avian-origin viruses, largely because of ɑ2,3-linked SIA availability on ciliated cells ^42,73,97,98^. Thus, it is probable that there are additional determinants of cell tropism in upper and lower respiratory epithelia than solely the presence of ɑ2,3-linked and ɑ2,6-linked SIAs, although glycan array analyses of contemporary H5 HA show a strong bias for avian-like SIAs ^35–37,41,56,57^. A more extensive analysis of the glycans present in human respiratory epithelial cell cultures, and in other tissues known to be infected by avian influenza viruses is needed to clearly delineate the potential glycans present on epithelial cells to help explain cell tropism differences.

Single amino acid mutations in avian-derived NS1 proteins have been shown to impact cytokine and chemokine responses, *in vitro* fitness, and pathogenicity ^99,100^. Among the NS1 effector-domain mutations, A/Washington/239/2024 NS1 N139D is notable because it has been associated with increased virulence of HPAI H5N6 in mice, and may contribute to mammalian adaptation ^101^. Residues 139 and 147 also overlap regions involved in NS1 nuclear export and localization ^102^. Collectively, these substitutions highlight candidate NS1 determinants that may affect host interaction, intracellular trafficking, or replication phenotypes. Despite the putative role of these NS1 mutations– as well as others previously mentioned in other gene segments– in host interactions, chemokine and cytokine responses to H5N1 and H1N1pdm09 infection were largely dictated by temperature and tissue type. There were lower levels of pro-inflammatory chemokines and cytokines that are essential to mounting functional antiviral responses during acute infection in both genotype B3.13 and D1.1 H5N1-infected hNECs and hBECs at 48 HPI, relative to those infected with the H1N1pdm09-like virus. These differences were the most distinct at 33°C at 48 HPI, and less pronounced at 37°C and 39°C, whereas at 96 HPI there were more significant differences within cultures incubated at 37°C and 39°C. hNECs tended to have more robust antiviral responses at both 48 HPI and 96 HPI, which might allude to different capacities for clearance between the upper and lower airway epithelia, though this area is understudied. Altogether, these data provide insight as to the status of mammalian adaptation among contemporary H5N1 viruses in physiologically relevant model systems of the human upper and lower respiratory tract and suggest that H5N1 genotypes B3.13 and D1.1 are not equally suited for replication at temperatures characteristic of the upper airways.

## ACKNOWLEDGEMENT

We thank the investigators at the US Centers for Disease Control for providing H5N1 isolates. This work was supported by the National Institutes of Health (NIH) contract 75N93021C00045 to the Johns Hopkins Centers of Excellence in Influenza Research and Response, NIH T32 AI007417, and the Richard Eliasberg Family Foundation. We thank the Pekosz laboratory for discussion of data and future directions We thank Maclaine Parish and Sabra Klein for use of the Luminex xMAP INTELLIFLEX System and acknowledge the MMI Microscopy Core at the Johns Hopkins Bloomberg School of Public Health for access to imaging instrumentation.

---

**Supplemental Table 1.** amino acid substitutions in A/H5N1 2.3.4.4b isolates (relative to A/Texas/37/2024)

| Isolate name | Genotype | Segment name |  |  |  |  |
| --- | --- | --- | --- | --- | --- | --- |
|  |  | PB1 | PA | NP | M | NS |
| A/Texas/37/2024 | B3.13 | -- | -- | -- | -- | -- |
| A/California/147/2024 | B3.13 | V392I | E142K, L219I, V432I, K497R | I119V | M1: S82N | NS1: R40Q, R67G |
| A/Washington/239/2024 | D1.1 | N16D, S59T, D75E, G154S, V171M, E172D, I179M, K207R, S376N, E390G, V392I, I568L, P587A, E614D, N694S | M61I, A85T, R113K, E142K, R269K, P277S, I322V, V323I, I348L, E351G, S388G, R391K, S400P, V441M, C489P, I545V, L558S, S608T, K626R | H52Y, N482S | M1: S82N, S85N, T87N, V200A, T227A | NS1: L7S, R40Q, E75G, S83P, P87S, S116C, N139D, L147I |

